# Rice BiP3 regulates immunity mediated by the PRRs XA3 and XA21 but not immunity mediated by the NB-LRR protein, Pi5

**DOI:** 10.1101/003533

**Authors:** Chang-Jin Park, Min-Young Song, Chi-Yeol Kim, Jong-Seong Jeon, Pamela C. Ronald

## Abstract

Plant innate immunity is mediated by pattern recognition receptors (PRRs) and intracellular NB-LRR (nucleotide-binding domain and leucine-rich repeat) proteins. Overexpression of the endoplasmic reticulum (ER) chaperone, luminal-binding protein 3 (BiP3) compromises resistance to *Xanthomonas oryzae* pv. *oryzae* (*Xoo*) mediated by the rice PRR XA21 (Park et al., PLoS ONE 5(2): e9262). Here we show that BiP3 overexpression also compromises resistance mediated by rice XA3, a PRR that provides broad-spectrum resistance to *Xoo*. In contrast, BiP3 overexpression has no effect on resistance mediated by rice Pi5, an NB-LRR protein that confers resistance to the fungal pathogen *Magnaporthe oryzae* (*M. oryzae*). Our results suggest that rice BiP3 regulates membrane-resident PRR-mediated immunity.

## INTRODUCTION

Plants recognize conserved microbial signatures via pattern recognition receptors (PRRs) (Ronald and Beutler 2010). Well-characterized PRRs include the Arabidopsis flagellin sensitive 2 (FLS2) receptor, the Arabidopsis elongation factor Tu receptor (EFR), and the rice XA21 (Xanthomonas resistance 21) receptor (Gomez-Gomez and Boller 2000; Song *et al*. 1995; Zipfel *et al*. 2006). These proteins share a similar structure: an extracellular leucine-rich repeat (LRR) domain, a transmembrane (TM) domain, and an intracellular non-arginine-aspartate (non-RD) kinase domain. The non-RD kinase motif is a hallmark of kinases involved in the initial stages of PRR-mediated immunity. In contrast, RD kinases are associated with a function in developmental processes (Dardick and Ronald 2006). Rice XA3 (also known as XA26) also shares this LRR-TM-non-RD kinase domain structure (Sun *et al*. 2004; Xiang *et al*. 2006). XA3 also confers another important trait characteristic of PRRs: broad-spectrum resistance. For these reasons, XA3 is predicted to be a PRR (Chen and Ronald 2011; Ronald and Beutler 2010; Schwessinger and Ronald 2012).

Membrane-bound PRRs are synthesized in the endoplasmic reticulum (ER) where they are subject to ER-quality control (Li *et al*. 2009; Nekrasov *et al*. 2009; Park *et al*. 2010; Saijo 2010). The ER-quality control is a conserved process in eukaryotic cells responsible for monitoring correct folding and processing of membrane and secretory proteins (Kleizen and Braakman 2004). Many ER proteins, including HSP70 luminal-binding protein (BiP), HSP40 ERdj3B, stromal-derived factor 2 (SDF2), calreticulin3 (CRT3), UDP-glucose glycoprotein glucosyl transferase (UGGT), and ER retention defective 2B (ERD2B), are involved in ER-quality control (Li *et al*. 2009; Liu and Howell 2010; Nekrasov *et al*. 2009). Recent research suggests that the ER-resident chaperone BiP, an integral protein of the ER quality control system, is one of the main chaperones regulating biogenesis and degradation of membrane-resident PRRs. For example, in BiP3-overexpressing rice plants, XA21-mediated immunity is compromised and accumulation/processing of XA21 after *Xoo* inoculation is significantly inhibited (Park *et al*. 2010). These results indicate that BiP3 regulates XA21 protein stability and processing and that this regulation is critical for resistance to *Xoo*. In Arabidopsis, a large ER chaperone complex BiP/ ERdj3B/SDF2 is required for the proper accumulation of EFR, further supporting a role for ER-quality control in membrane-resident PRR–mediated function (Nekrasov *et al*. 2009).

In contrast to PRRs that recognize conserved microbial signatures, intracellular NB-LRR proteins, containing nucleotide-binding and leucine-rich repeat domains, perceive highly variable pathogen-derived effectors in the cytoplasm directly or indirectly. Many NB-LRR proteins, including Arabidopsis RPM1, RPS2, and RPS5, tobacco N and rice Pi-ta and Pi5, have been characterized (Bent *et al*. 1994; Grant *et al*. 1995; Jeon *et al*. 2003; Jia *et al*. 2000; Warren *et al*. 1998). In tobacco, a number of ER resident chaperones, including BiP, are up-regulated during the earlier stages of N immune receptor-mediated defense against *Tobacco mosaic virus* (Caplan *et al*. 2009). In Arabidopsis, BiP2 is also involved in folding and secretion of pathogenesis-related (PR) proteins (Wang *et al*. 2005). In a *bip2* mutant, increased PR protein synthesis after benzothiadiazole S-methylester (BTH, salicylic acid analog) treatment is not accompanied by a concomitant increase in BiP protein accumulation, resulting in impaired BTH-induced resistance. These results suggest that components of ER-quality control may also play a role in NB-LRR-mediated immunity (Padmanabhan and Dinesh-Kumar 2010).

A differential requirement for gene governing ER-quality control has been observed in structurally-related receptors. For example, Arabidopsis FLS2-mediated responses are not impaired in the mutants, *sdf2*, *crt3*, *uggt*, and *erd2b* (Li *et al*. 2009; Nekrasov *et al*. 2009). In contrast, these genes which encode ER proteins, are all required for EFR-mediated signal transduction. Similarly, overexpression of BiP3 in rice does not affect rice brassinosteroid-insensitive 1 (OsBRI1)-mediated signaling, even though OsBRI1 shows an overall structural similarity with XA21. In contrast to XA21, OsBRI1 carries the RD class of kinases and regulates growth and developmental responses. As observed for rice, Arabidopsis BiPs fail to interact with wild-type BRI1 (Hong *et al*. 2008). In animals, processing of Toll-like receptors (TLRs), which are key determinants of the innate immune response, also require specific ER chaperones. For example, despite of its role as a general housekeeping chaperone, mouse ER gp96 is specific for processing TLR2, 4, 5, 7, and 9 in macrophages (Liu *et al*. 2010; Takahashi *et al*. 2007; Yang *et al*. 2007). Although gp96-deficient macrophages fail to respond to flagellin, the ligand for TLR5, mutant macrophages display normal development and activation by interferon-c, tumor necrosis factor-a, and interleukin-1b (Yang *et al*. 2007).

Based on these results, we hypothesize that ER chaperones are specific to their substrates, differentially regulating plant responses. To test this hypothesis, we chose two different types of receptors, the membrane-resident PRR XA3 and the intracellular NB-LRR protein Pi5, and assessed if BiP3 overexpression affects these immune responses. We found that overexpression of BiP3 compromises XA3-mediated immunity to *Xanthomonas oryzae* pv. *oryzae* (*Xoo*) but not Pi5-mediated immunity to *Magnaporthe oryzae* (*M. oryzae*). These results indicate that BiP3 regulates PRRs and that BiP3 overexpression does not lead to a general ER stress response.

## MATERIAL AND METHODS

### Plant materials and growth conditions

The endoplasmic reticulum (ER) chaperone *BiP3* overexpressing (BiP3ox) transgenic line 1A-6 (Park *et al*. 2010) and its background cultivar Kitaake, IRBB3 monogenic line carrying *Xa3* (Xiang *et al*. 2006), and IRBL5-M monogenic line carrying *Pi5* (Yi *et al*. 2004) were used in this study. The IRBB3 and IRBL5-M lines were crossed with BiP3ox 1A-6 to generate BiP3ox/XA3 and BiP3ox/Pi5 plants, respectively. Self-pollinated seeds (F_2_) were collected and were used in pathogen inoculations. Rice plants were grown in a greenhouse at 30°C during the day and at 20°C at night in a light/dark cycle of 14/10 hours.

### *Xanthomonas oryzae* pv. *oryzae* inoculation and lesion length measurements

Rice plants were grown in the greenhouse normally until they were six-week-old and transferred to the growth chamber. Growth chambers were set on 14 hours light:10 hours dark photoperiod, 28/26°C temperature cycle, and 85/90% humidity. The *Xoo* strains PXO61 and PXO79 were used to inoculate rice by the scissors-dip method (Chern *et al*. 2005; Song *et al*. 1995). *Xoo* strains PXO61 and PXO79 were grown on PSA plate (Peptone Sucrose Agar, 10 g/L peptone, 10 g/L sucrose, 1 g/L glutamic acid, 16 g/L agar, and pH 7.0) containing cephalexin (100 µg/L) for three days and suspended with water at OD=0.5 (600 nm) for inoculation. Only the top two to three expanded leaves of each tiller were inoculated.

### *Magnaporthe oryzae* inoculation and disease evaluation

*M. oryzae* R01-1, which is incompatible with the *Pi5*-carrying line (Lee *et al*. 2009) and compatible with the cultivar Kitaake, has been commonly used to evaluate disease resistance. *M. oryzae* R01-1 was grown on rice flour medium containing 20 g/L rice flour powder, 10 g/L dextrose, and 12 g/L agar in the dark for 2 weeks at 22°C (Lee *et al*. 2009). Conidia were induced for 5 days by scratching the plate surface with a sterilized loop. Agar blocks covered with spores were placed on the injured spot of 2.0 mm diameter according to the leaf punch inoculation (Takahashi *et al*. 1999). The second fully expanded leaves from the top of five-week-old plants were used for *M. oryzae* inoculation. The inoculated plants were placed in sealed containers to maintain humidity in the dark for 24 hours at 24°C and then incubated at the same relative humidity under a 14/10 hours (light/dark) photoperiod. For disease evaluation, blast lesion lengths were measured 10 days after inoculation.

### Genomic DNA isolation and PCR analysis

Genomic DNAs were extracted from leaves of individual plants following the protocol described previously (Chen and Ronald 1999). The gene-specific primers for PCR were as follows: for *BiP3*, 5**′**-GCTGCTGCTATTGCGTACGGTTTGGACA-3**′** and 5**′**-AATCATCGCAAGACCGGCAACAGG-3**′**; for *Pi5-1*, 5**′**-TTATGAGATTAGGAGTGTAT-3**′** and 5**′**-ATGTAAAGGCAAAAGCTGAT-3**′**; and for *Pi5-2*, 5**′**-CTCTTGGTGATCTTTGTTAC-3**′** and 5**′**-GGATGATGTGATCTGCAGAG-3**′**; and for *Xa3*, 5**′-** CACCCCACGCAAGCCTCTCA-3**′** and 5**′-**CTCCGTCATCAGCCATACACTCAC-3**′**. We carried out PCR experiments. PCR conditions were pre-denaturation for 3 min at 94°C, followed by 35 cycles of polymerization, each consisting of 30 s of denaturation at 94°C, annealing for 30 s at 58°C, and an extension step of 30 s at 72°C. The PCR product was separated on a 1.5% agarose gel and stained with ethidium bromide to detect the amplicons.

## 002501v1

### Overexpression of *BiP3* compromises membrane-resident PRR XA3-mediated immunity

As is typical of PRRs, XA3 confers broad-spectrum resistance to *Xoo* strains including PXO61 and PXO79 (Cao *et al*. 2007). To explore the biological role of BiP3 in XA3-mediated immunity, we first investigated the presence of a functional *Xa3* gene in the genome of rice cultivar Kitaake. Using the scissors-dip-method, we inoculated Kitaake plants with *Xoo* strains PXO61 and PXO79. Figure 1A shows typical leaves from Kitaake wild type and *Xa3*-possessing IRBB3 plants 14 days after inoculation with *Xoo* strains. While IRBB3 was highly resistant, showing short lesions (approximately 4 to 5 cm), the inoculated leaves of Kitaake plants developed long water-soaked lesions (approximately 17 to 20 cm) typical of bacterial blight disease (Fig. 1B). These results indicate that Kitaake plants do not possess a functional *Xa3* gene.

**Fig. 1.**
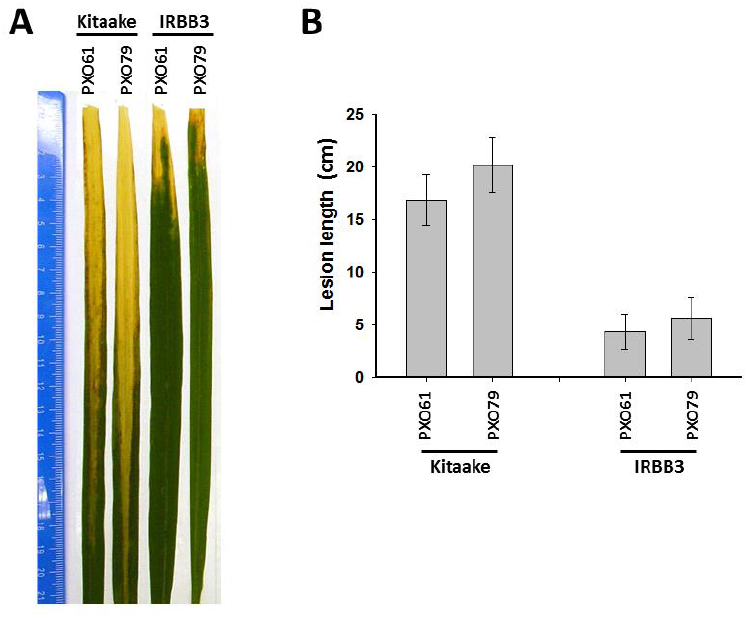
Rice cultivar Kitaake does not carry a functional *Xa3*. (A) Kitaake plants were susceptible to *Xoo* strains PXO61 and PXO79. Kitaake and IRBB3 plants were inoculated with *Xoo* strains PXO61 or PXO79 at six-week-old age and a representative leaf was harvested 14 days after inoculation for photograph. (B) Kitaake plants developed long lesion lengths after inoculation with *Xoo* strains PXO61 or PXO79. Lesion lengths of Kitaake and IRBB3 plants were measured 14 days after inoculation. Each data point represents the average and standard deviation of at least three samples.

To generate XA3 plants overexpressing *BiP3*, we first crossed IRBB3 (XA3, pollen donor) with transgenic Kitaake overexpressing *BiP3* (BiP3ox 1A-6, pollen recipient) (Park *et al*. 2010). The presence of the *BiP3ox* and/or *Xa3* genes in the resulting F_1_ was confirmed by PCR analysis with primers specific for *BiP3ox* or *Xa3* (data not shown). The F_2_ segregating progenies (a total of 41 segregating progeny) were inoculated with *Xoo* strain PXO61, and examined for cosegregation of the genotype with lesion length developments (Fig. 2A). XA3 plants lacking *BiP3ox* (BiP3ox/XA3, -/+) developed short lesion lengths (approximately 2 to 3 cm), indicating that *Xa3* confers resistance to *Xoo* strain PXO61. In contrast, all XA3 plants carrying *BiP3ox* (BiP3ox/XA3, +/+) displayed significantly compromised resistance to *Xoo* strain PXO61. The long lesions formed (approximately 15 cm) are similar to those of the segregants lacking *Xa3* (BiP3ox/XA3, -/- and +/-). Bacterial growth was also monitored over time (Fig. 2B). *Xoo* strain PXO61 populations in XA3 plants (BiP3ox/XA3, -/+) reached approximately 5.5×10^7^ colony-forming units per leaf (cfu/leaf), whereas the population in the segregants lacking *Xa3* (BiP3ox/XA3, -/- and +/-) reached to more than 9.1×10^8^ cfu/leaf. In plants carrying both *Xa3* and *BiP3ox* (BiP3ox/XA3, +/+), the bacterial populations grew to 8.3×10^8^ cfu/leaf, a greater than fifteen-fold increase compared to the XA3 plants. The F_2_ progenies inoculated with another *Xoo* strain PXO79 showed similar results with increased lesion length developments and bacterial populations (Fig. 2C and D). These results demonstrate that overexpression of BiP3 compromises XA3-mediated immunity against *Xoo*.

**Fig. 2.**
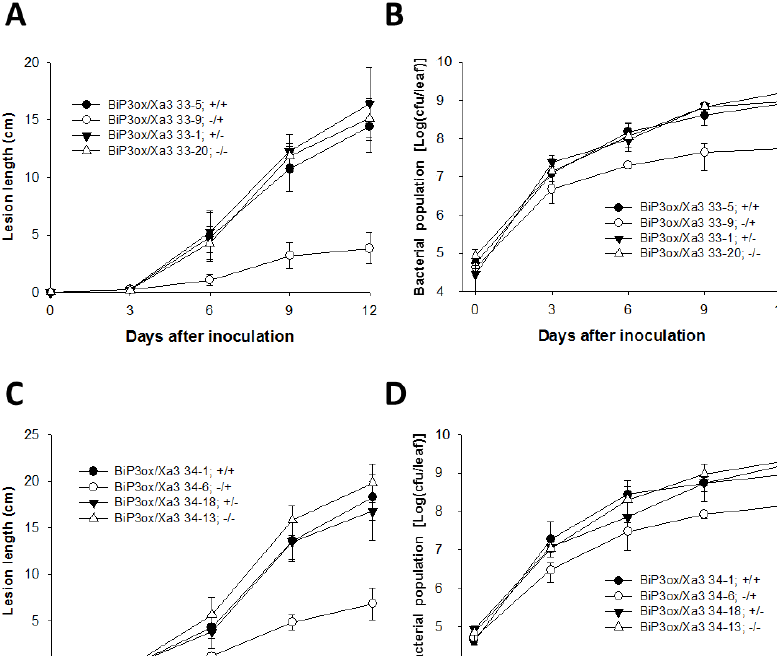
Overexpression of *BiP3* compromises XA3-mediated resistance. (A) Lesion length developments of *Xoo* PXO61-inoculated segregants (F_2_) of BiP3ox/XA3. Each data point represents the average and standard deviation of at least three samples. BiP3ox/XA3 (*BiP3ox*/*Xa3*, +/+); BiP3ox (*BiP3ox*/*Xa3*, +/-); XA3 (*BiP3ox*/*Xa3*, -/+); Null segregants (*BiP3ox*/*Xa3*, -/-). (B) *Xoo* PXO61 populations in segregants (F_2_) of BiP3ox/XA3. For each time point, the bacterial populations were separately determined for three leaves. Capped, vertical bars represent standard deviation of values (cfu/leaf) from three samples. Experiments were repeated over two times with similar results. BiP3ox/XA3 (*BiP3ox*/*Xa3*, +/+); BiP3ox (*BiP3ox*/*Xa3*, +/-); XA3 (*BiP3ox*/*Xa3*, -/+); Null segregants (*BiP3ox*/*Xa3*, -/-). (C) and (D) Lesion length developments and bacterial populations after *Xoo* strain PXO79 inoculation.

### Overexpressed *BiP3* does not affect Pi5-mediated immunity

We have previously shown that Kitaake plants overexpressing *BiP3*, lacking *Xa21*, do not show alterations in resistance to *Xoo* strain PXO99, suggesting that an increased level of BiP3 does not affect the general defense response to bacterial pathogen *Xoo* (Park *et al*. 2010). To determine if enhanced expression of *BiP3* can affect the resistance level to fungal pathogen, we inoculated the BiP3ox and the Kitaake wild type plants with *M. oryzae* R01-1. Figure 3A shows a typical leaf from each of the inoculated rice plants 10 days after *M. oryzae* inoculation. We found both BiP3ox and Kitaake plants displayed similar blast lesions of approximately 1.0 to 1.2 cm (Fig. 3B), indicating that *BiP3* overexpression does not affect general resistance to *M. oryzae*.

**Fig. 3.**
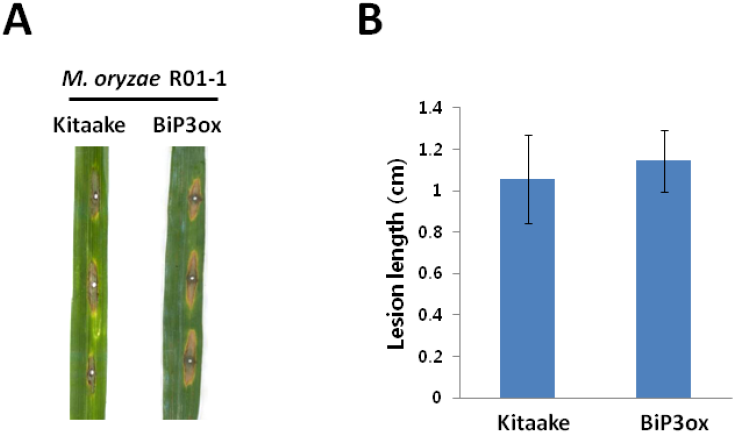
BiP3 overexpression does not affect the general defense response to *M. oryzae*. (A) Image of disease lesions on one leaf from Kitaake and BiP3ox 10 days after *M. oryzae* R01-1 inoculation. Plants were inoculated at five-week-old age and harvested for photograph. (B) Lesion lengths of Kitaake and BiP3ox were measure 10 days after inoculation. Each data point represents the average and standard deviation of at least 30 samples.

To further determine if enhanced expression of *BiP3* can affect NB-LRR-mediated resistance to *M. oryzae*, we examined the effect of *BiP3* overexpression on *Pi5*-mediated resistance to *M. oryzae*. *Pi5* resistance is controlled by two genes, *Pi5-1* and *Pi5-2*, which encode coiled-coil (CC)-NB-LRR proteins (Lee *et al*. 2009). We produced double transgenic plants carrying both *Pi5* and *BiP3ox* by crossing the IRBL5-M carrying *Pi5* (pollen donor) and BiP3ox 1A-6 lines (pollen recipient). The presence of the *BiP3ox* and/or *Pi5* in the resulting F_1_ was confirmed by the PCR analysis with primers specific for *BiP3ox* or *Pi5* (data not shown). The F_2_ segregating individual plants (a total of 64 segregating progeny) from 5 F_1_ lines (*BiP3ox*/*Pi5*, +/+) were then inoculated with *M. oryzae* R01-1 (Fig. 4A and B). We found that all BiP3ox/Pi5 plants carrying both genes (+/+; #2, 4, 5, 6, 7, 8, and 10 in Fig. 4A) remained completely resistant to the *M. oryzae* strain comparable to the parental line IRBL5-M or segregants carrying *Pi5* but lacking BiP3ox (BiP3ox/Pi5, -/+; #9 in Fig. 4A). In contrast, the parental BiP3ox line or segregants carrying *BiP3ox* but lacking *Pi5* (BiP3ox/Pi5, +/-; #1 and 3 in Fig. 4A) were all susceptible to the *M. oryzae*. These results indicate that overexpression of *BiP3* does not interfere with either general or *Pi5*-mediated resistance to *M. oryzae*.

**Fig. 4.**
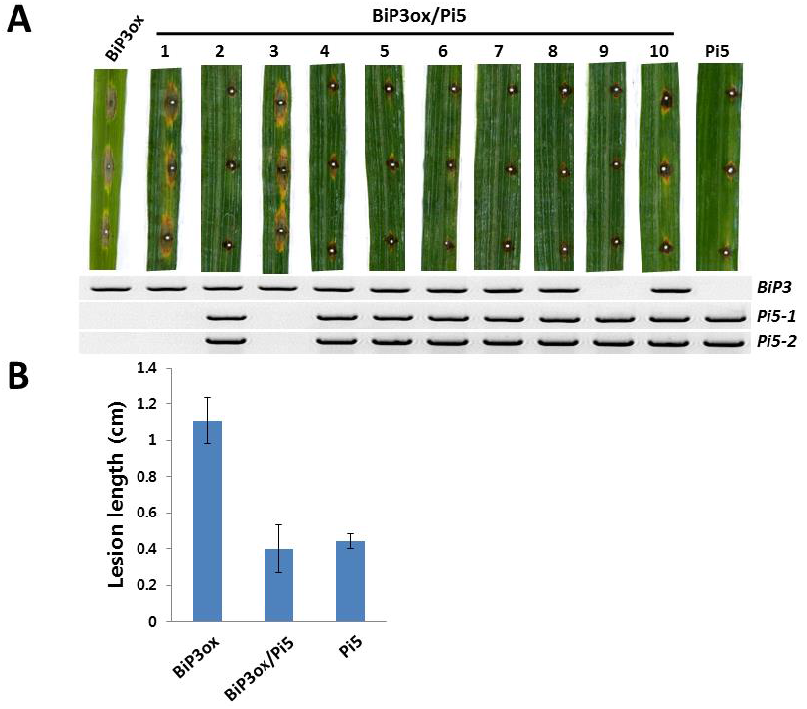
Overexpression of BiP3 does not compromise Pi5-mediated resistance. (A) Phenotype/genotype cosegregation analysis of a total of 64 segregating F_2_ progeny [*BiP3ox*/*Pi5*; 43 (+/+), 6 (-/+), 13 (+/-), 2 (-/-)] derived from BiP3ox/Pi5 F_1_ line in response to *M. oryzae* R01-1 inoculation. Detailed inoculation and genotyping methods are described in the materials and methods. (B) Lesion lengths of F_2_ progeny derived from BiP3ox/Pi5 F_1_ line in response to *M. oryzae* R01-1 inoculation. Each data point represents the average and standard deviation of at least 30 samples. BiP3ox (*BiP3ox*/*Pi5*, +/-); BiP3ox/Pi5 (*BiP3ox*/*Pi5*, +/+); Pi5 (*BiP3ox*/*Pi5*, -/+).

## DISCUSSION

After PRRs are synthesized in the ER, the folding states of the PRRs are monitored by ER chaperones. BiP is one of the most abundant chaperones, functioning in the ER-quality control (Liu and Howell 2010; Otero *et al*. 2010). In Arabidopsis, the ER quality control complex SDF2/ERdj3B/BiP is required for EFR accumulation and the EFR-mediated immune response (Li *et al*. 2009; Saijo 2010). In contrast, in rice, *BiP3* overexpression significantly inhibits XA21 accumulation. These results suggest that overexpression of BiP3 results in XA21 degradation possibly via ER-associated degradation as an ER-quality control mechanism (Park *et al*. 2010). The genetic requirement for BiP in Arabidopsis PRR-mediated immune responses has been difficult to elucidate because single mutants (*bip1*, *bip2*, or *bip3*) and a double mutant (*bip1 bip2*) all display normal EFR-mediated immune response (Nekrasov *et al*. 2009). Silencing of all three Arabidopsis *BiP* genes caused lethality (Hong *et al*. 2008). Therefore, the direct involvement of Arabidopsis BiPs in the biogenesis of EFR has not been demonstrated (Li *et al*. 2009). Similarly in rice, mutants silenced for *BiP3*, display no obvious phenotypes (Park *et al*. 2010). However, we have demonstrated that BiP3 directly interacts with XA21 in the ER and that transgenic plants overexpressing BiP3 are compromised in XA21-mediated immune responses to *Xoo,* highlighting the important role of *BiP3* in the XA21-mediated immune response (Park *et al*. 2010). Here, we also show that overexpression of BiP3 compromises resistance mediated by the PRR XA3.

The rapid transcriptional induction and protein accumulation of ER chaperones including BiP in response to various pathogens and/or effector treatments (Caplan *et al*. 2009; Foster-Hartnett *et al*. 2007; Fujiwara *et al*. 2006; Jelitto-Van Dooren *et al*. 1999; Zulak *et al*. 2009) suggests that BiP may also play a role in NB-LRR protein-mediated resistance. Supporting this hypothesis, confocal microscopy experiments examining Arabidopsis NB-LRR proteins found many of NB-LRRs to be localized to the ER (Caplan *et al*. 2009). Thus, the ER-resident NB-LRR proteins likely require ER chaperones for maturation (Caplan *et al*. 2009). Here we show that rice BiP3 overexpression does not affect Pi5-mediated resistance to *M. oryzae.* These results suggest that overexpressed BiP3 does not have a role in the biogenesis of Pi5 itself or in Pi5-mediated signaling. These data also indicate that BiP3 overexpression does not lead to a general ER stress response affecting resistance. Thus it is likely that other ER chaperones are involved in the Pi5-mediated immune response to *M. oryzae*.

This research has focused on BiP’s involvement in immunity mediated by the PRR XA3 and the NB-LRR Pi5 protein. Rice plants overexpressing *BiP3* are compromised in membrane-resident PRR XA3-mediated immunity to *Xoo* but not intracellular NB-LRR Pi5-mediated immunity to *M. oryzae*. These results suggest that BiP3 is involved in the biogenesis of other PRRs, rather than a general role in the resistance caused by non-specific ER stress.

## ACKNOWLEDGMENTS

We thank Dr. Benjamin Schwessinger and Dr. Rita Sharma for critical reading of the manuscript. This work was, in part, supported by grants from the National Science Foundation (NSF, IOS-0817738 to P.C.R.), the Next-Generation BioGreen 21 Program, Rural Development Administration, the Korean Ministry of Agriculture, Food and Rural Affairs (PJ008156 to J.S.J), and the Basic Science Research Programs through the National Research Foundation of Korea (NRF) by the Ministry of Science, ICT & Future Planing (2013R1A1A1059051 to C.J.P) and by the Ministry of Education (2013R1A1A2061860 to M.Y.S).

## Authors’ contributions

C.J.P., J.S.J., and P.C.R. contributed to project design. C.J.P., M.Y.S., and C.Y.K. performed experiments. C. J. P., J.S.J., and P.C.R. wrote the paper. All authors discussed the results and commented on the manuscript.

